# Genomic disruption of type II vitellogenin leads to vitellin membrane deficiencies and significant mortalities at early stages of embryonic development in zebrafish (*Danio rerio*)

**DOI:** 10.1101/2023.07.24.550261

**Authors:** Ozlem Yilmaz, Emmanuelle Com, Charles Pineau, Julien Bobe

**Affiliations:** INRA, UR1037, Laboratory of Fish Physiology and Genomics, Campus de Beaulieu, 35042 Rennes Cedex, France; Univ Rennes, Inserm, EHESP, Irset (Institut de Recherche en Santé, Environnement et Travail) - UMR_S 1085, F-35000 Rennes, France; Univ Rennes, CNRS, Inserm, Biosit UAR 3480 US_S 018, Protim core facility, F-35000 Rennes, France; Institute of Marine Research, Austevoll Research Station, 5392, Storebø, Norway

**Keywords:** *CRISPR/Cas9*, *knock out*, *vitellogenins*, *zebrafish*

## Abstract

Type II vitellogenin (Vtg2), the second most abundant type of vitellogenin in zebrafish eggs, is a major source of nutrients for early embryonic development. The main objective of this study was to determine the specific functions and essentiality of *vtg2* in zebrafish early development using CRISPR/Cas9 genome editing tool. A 2811 bp deletion on gDNA was detected in *vtg2*-mutant zebrafish via PCR genotyping and sequencing. Introduced mutation caused vitelline membrane deficiencies and significant mortalities of mutant offspring. Further effects on female fecundity, egg fertilization rate and *vtg2* gene expression and Vtgs abundance in liver were observed in F2, while effects on embryo hatching, survival rates, proteomic profiles and abundances were observed in F3 generations of the *vtg2*-mutant line. No change in *vtg2* transcript has been detected, however, Vtg2 abundance in F2 female liver was 5x, and in 1 hpf F3 *vtg2*-mutant embryos was 3.8x less than Wt (*p* < 0.05). All Vtgs, except Vtg7, declined in abundance in 1 hpf F3 *vtg2*-mutant embryos (*p* < 0.05). Fecundity was unaffected while fertilization rate was more than halved in F2 *vtg2*-mutant females (*p* < 0.05). Hatching rate was significantly higher in F3 *vtg2*-mutant embryos in comparison to Wt embryos. Survival rate declined drastically to 29 % and 18 % at 24 hpf and 20 dpf, respectively, in F3 *vtg2*-mutant embryos. Pericardial, yolk sac/abdominal edema and spinal lordosis were evident at later stages in the surviving F3 *vtg*2*-*mutant larvae. Overrepresentation and high expression of histones, zona pellucida proteins, lectins, and protein degradation related proteins in F3 *vtg2*-mutant embryos provide evidence to impaired mechanisms involved in vitellin membrane formation. Findings of this study imply a potential function of Vtg2 in acquisition of vitellin membrane integrity, among other reproductive functions, and therefore, its essentiality in early zebrafish embryo development.

**AUTHOR SUMMARY:** Vitellogenins (Vtgs) are major yolk nutrient precursors supporting early vertebrate development. Most species have multiple forms of Vtg, but little is known about their individual roles in reproduction and it is uncertain which forms are essential for successful development or at what stage(s) of development they are required. This study employed a CRISPR/Cas9 gene knock out (KO) to assess essentiality and functionality of Vtg2 in zebrafish, in continuation to a previously published work on type I and type III *vtgs* KO. The findings of this study, in combination with the previous findings, present a new model of Vtg functionality. Accordingly, Vtg2 contribute to regulation of fecundity and fertilization in female reproduction while make essential contributions to embryonic morphogenesis, hatching and embryonic and larval kinetics and survival. In addition, Vtg2 is critically important to proper formation of the vitellin membrane, and thus, to embryogenesis and later development. Our novel findings provide, for the first time, empirical evidence that the Vtg2 are essential, having critical requisite functions during oogenesis and embryonic and larval development. The overall results substantiate the concept that each type of Vtg is specialized to play unique roles in reproduction and development.

## 1. INTRODUCTION

Vitellogenin 2 is part of the complex zebrafish (*Danio rerio*) multiple Vtgs system which contains five type-I Vtgs (Vtg 1, 4, 5, 6 and 7), two type-II Vtgs (Vtg2 and Vtg8), and one type-III Vtg (Vtg3). These three types correspond to the spiny-rayed teleost (Acanthomorpha) Vtgs; VtgAa (type-I Vtgs), VtgAb (type-II Vtgs) and VtgC (type-III Vtgs). The linear yolk protein domain structure of complete teleost Vtgs is: NH_2_-lipovitellin heavy chain (LvH)-phosvitin (Pv)-lipovitellin light chain (LvL)-beta component (β’c)-C-terminal component (Ct)-COOH [1, 2]. The type I zebrafish Vtgs and the type III Vtg (Vtg3) are incomplete lacking β′-c and Ct domains and both Pv and the β’-c and Ct domains, respectively, while the type II Vtgs are complete forms containing all yolk protein domains [3].

The roles that different types of Vtg play in oocyte growth and maturation and in embryonic and larval development has been the target of attention for decades [2, 4]. Yolk proteins derived from VtgAa are reported to be highly susceptible to proteolytic degradation by cathepsins during oocyte maturation, yielding a pool of free amino acids (FAA) that osmotically assist oocyte hydration and acquisition of proper egg buoyancy [5, 6]), and that also serve as critical nutrients during early embryogenesis of some marine species spawning pelagic eggs [7, 8]. The major yolk protein derived from the corresponding VtgAb (LvHAb) is less susceptible to maturational proteolysis.

In zebrafish, a freshwater fish spawning demersal eggs, however, it has been demonstrated that type I Vtgs, the major contributors to yolk proteins in zebrafish eggs, are essential for early embryonic and late larval development while type III Vtg, made of Lv domains only and being the least abundant form of Vtg, is a critical resource for early embryonic development [9]. These two types of Vtgs have been proven to be essentially required for the developmental competence of zebrafish eggs and/or offspring [9]. However, the function of type VtgII in early zebrafish development remained to be investigated. Therefore, the main objective of this study was to discover whether type-II Vtgs are required for zebrafish reproduction, and to identify specific developmental periods and processes to which they significantly contribute, by investigating the effects of knock out (KO) of their respective genes using the CRISPR/Cas9 gene-editing tool.

## 2. RESULTS

A large deletion mutation of 2811 bp of gDNA was introduced into zebrafish *vtg2* via CRISPR/Cas9 genome editing. Schematic representation of the general strategy for CRISPR target design and application is given in **Fig 1A-C**. The introduced deletion involved 1692 bp of the *vtg2* transcript, encoding 564 aa of its respective protein and resulted in a frame shift on the ORF (**Fig 1**, **Fig 2, S1 Fig**). This was expected to alter the structure of the deduced LvH chain and disturb the function of the remaining domains (Pv, LvL, Bc, Ct) (**Fig 2, S1 Fig**). The introduced mutation was detected by genotyping via conventional PCR screening of gDNA at each generation using combinations of primers flanking each altered target site (**Fig 1C**).

**Fig 1.**
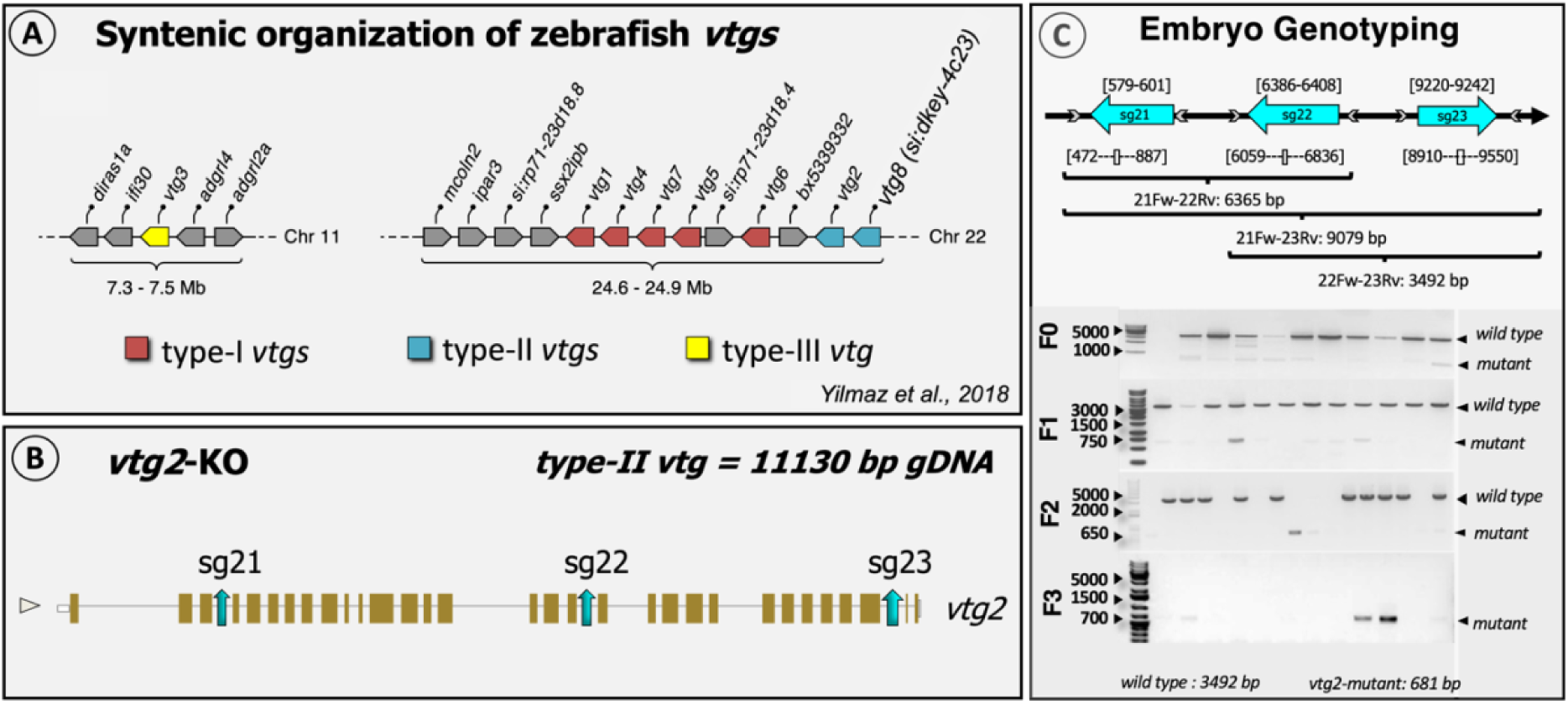
Schematic representation of the general strategy for CRISPR target design. **A.** Synthetic organization of *vtg* genes on zebrafish genome (from [3]). **B.** Target locations on zebrafish *vtg2* genomic structure. Target sites are shown by blue colored arrows labeled as sg followed by 1, 2 or 3 indicating the targeted zebrafish *vtg* type and the number of the target site (*i.e*., sg21, sg22, and sg23: single guide RNAs (sgRNAs) for target sites 1, 2, and 3 for *vtg2*, respectively. **C.** Detection of CRISPR/Cas9- introduced mutation by embryo genotyping. Target sites are shown in top left panel by blue colored arrows labeled as sg followed by 1, 2 or 3 indicating the targeted zebrafish *vtg* type and the number of the target site (*i.e.,* sg21, sg22, and sg23: single guide RNAs (sgRNAs) for target sites 1, 2, and 3 for *vtg2*, respectively. Arrows are oriented to indicate the sense/antisense orientation of each target. Numbers above each target site specify its exact location by nucleotide in the genomic sequence of the zebrafish *vtg2*. Primers used in screening for introduced mutations by PCR are shown as grey arrowheads which are oriented to indicate the sense/antisense orientation of the primer. Numbers below each primer site indicate its exact position by nucleotide in the genomic sequence of the targeted gene (see also **S1 Fig**). Horizontal brackets below indicate areas screened for mutations by PCR using selected primer combinations; text below the brackets indicates the primer pair followed by the size of the band (bp) expected for wild type gDNA in agarose gel electrophoresis. 21Fw, *vtg2* target1 forward primer; 22Rv, *vtg2* target2 reverse primer; 23Rv, *vtg2* target3 reverse primer; 22Fw, *vtg2* target2 forward primer. Primer sequences are given in **S3 Table**. Bottom left panel illustrate genotyping of embryos at 24 h post-fertilization (hpf) by PCR for *vtg2*-mutant line, from the F0 to F3 generation. F0 indicates the generation reared from microinjected embryos and F1-3 represent offspring raised from each subsequent generation. The agarose gel electrophoresis results shown here represent screening of 10-17 randomly sampled embryos as representatives of their generations. Bands comprised of wild type intact gDNA (3492 bp) and mutated gDNA (681 bp) are shown and highlighted by black arrowheads on the right side of each panel.

**Fig 2.**
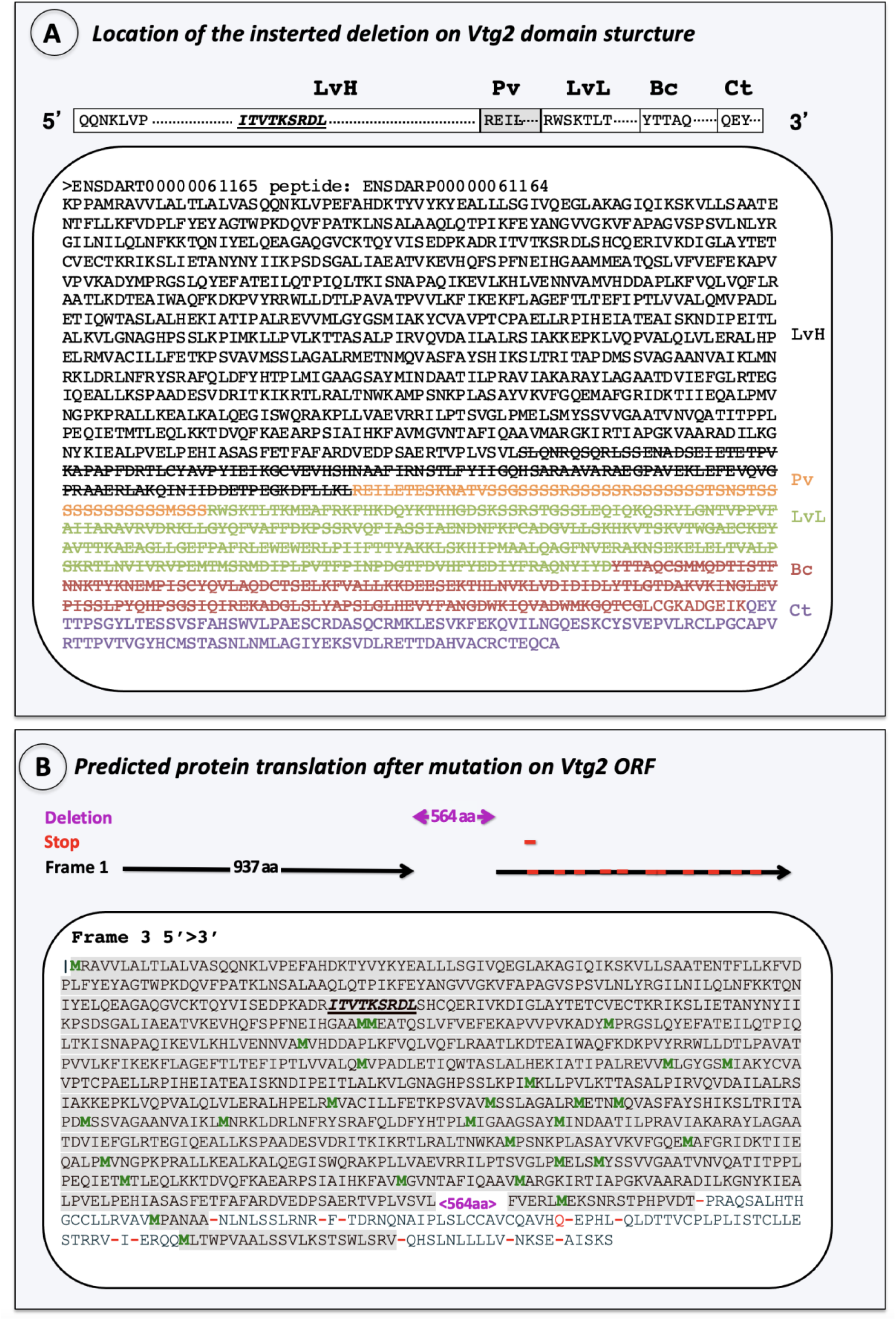
Characterization of the introduced mutation. A. Location on Vtg2 polypeptide structure. The yolk protein domain structures of Vtg2 is pictured in 5’ > 3’ orientation above panel. Horizontal bars represent the heavy and light chain lipovitellins (LvH, LvL), phosvitin (Pv), beta component (Bc) and C-terminal (Ct) domains of the Vtg2 and are labeled above in large bold type. Sequences within these bars indicate the N-terminus of each yolk protein domain, the starting points of which are also indicated by shade color change to the polypeptide sequence shown below. Each domain is labeled to the right with the corresponding shade color. LvH: black, Pv: orange, LvL: light green, Bc: red, Ct: purple. Cas9 created mutations (large deletions) are indicated with stroked letters representing amino acid (aa) residues. **B.** Frame shift caused by the introduced mutation on the Vtg2 open reading frame. A total of 973 aa which were not altered by the introduced mutation are indicated in grey shaded letters. Introduced mutation of 564 aa is indicated in magenta font set (< 564 aa >). Methionine residues are shown in green font set and “stop codons” are indicated by red dashes. Receptor binding site is indicated by bold, italic and underlined aa motif.

Efficacy of CRISPR/Cas9 was ∼54 %, however, mutation transmission to subsequent generations was extremely low (0.25 %) in this study. Only a single F0 male zebrafish exhibiting a mosaic mutant double band pattern was detected and used as the founder of F1, and only two heterozygous F1 male zebrafish were detected and crossed with wild type females to produce F2 generations. At this level, mutant F2 generations were expected to be heterozygous with a single mutated *vtg2* allele (Ht; *vtg2* -/+). Therefore, all individuals revealed a genotype with the mutation band pattern (with or without the wild type band) were considered as heterozygous (Ht; *vtg2* -/+) mutant and referred to as *vtg2*-mutant in this study. Ultimately, F3 offspring batches that were subject to proteomic and phenotyping analyses were a mix of homozygous (Hm; *vtg2* -/-), heterozygous (Ht; *vtg2* -/+) and wild type (Wt; *vtg2* +/+) individuals with half maternal contribution because their female parent is a heterozygous with a single *vtg2* allele (Ht; *vtg2* -/+) (**Fig 3**).

**Fig 3.**
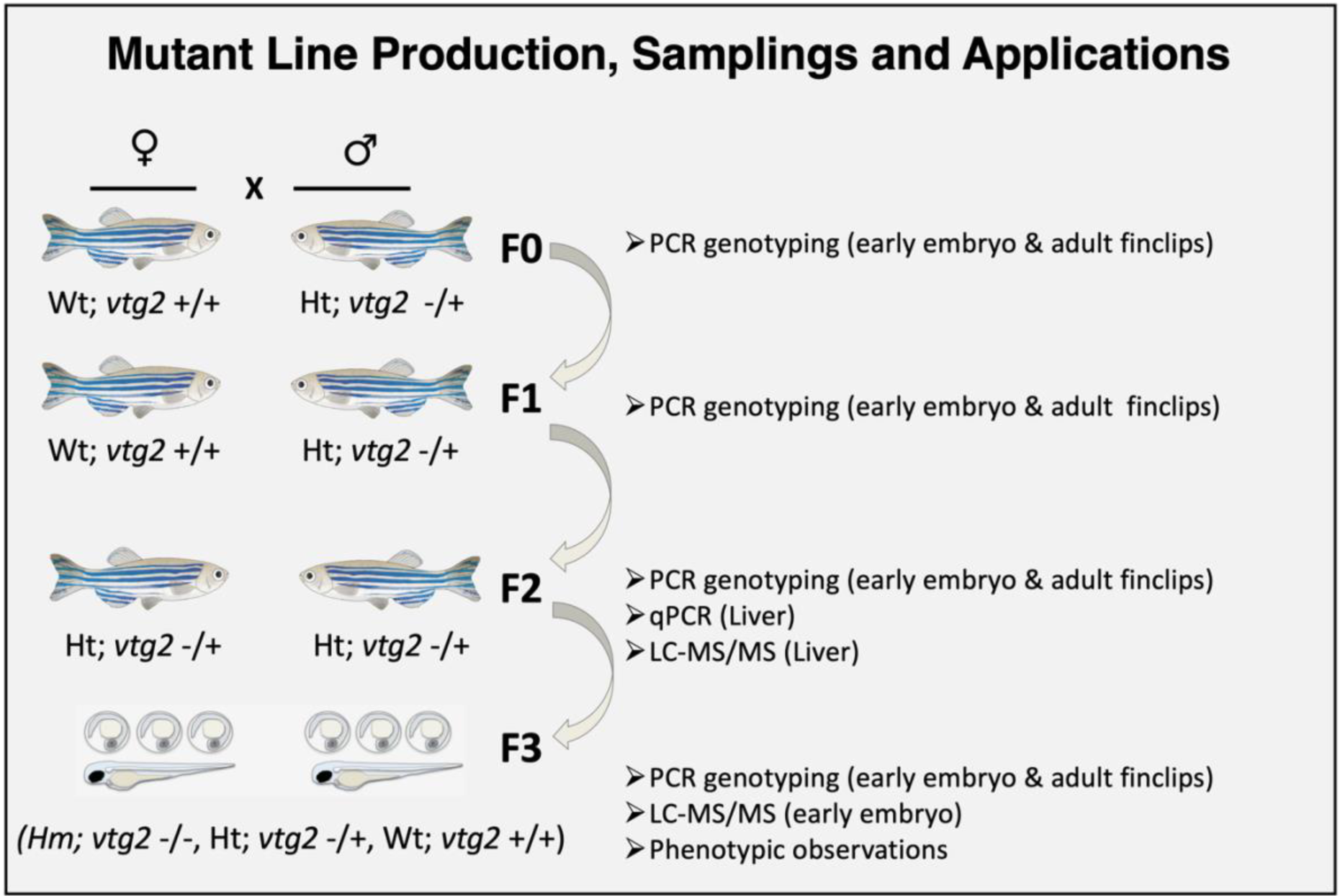
Production of F3 generation *vtg2*-mutants and samplings. The general strategy followed to establish zebrafish *vtg2*-mutant lines is shown above. This process involved stepwise reproductive crosses (indicated by X) between males (♂) and females (♀) indicated here with zebrafish icons. F0-3 represents the zebrafish generations produced in the process. Biological samples and the applications for which they were used are indicated on the right side of the panel. Wt; *vtg2* +/+, wild type fish with two *vtg2* alleles, Ht; *vtg2* -/+, heterozygous fish carrying only one *vtg2* allele, Hm; *vtg2* -/-, homozygous fish lacking both *vtg2* alleles.

The relative expression of *vtg2* transcript in F2 generation *vtg2*-mutant female (Ht; *vtg2* -/+) liver did not differ from those of Wt female liver (p > 0.05). In contrast, the relative abundance of individual Vtgs in liver of F2 *vtg2*-mutant females (*vtg2*-mutant; 11,6275 ± 3,31, Wt; 2,31 ± 0,65), as well as in their 1 hpf F3 *vtg2*-mutant embryos (*vtg2*-mutant; 44,405 ± 2,46, Wt; 169,7675 ± 12,1), was significantly reduced in comparison to those of Wt (*p* < 0.05) (**Fig 4**).

**Fig 4.**
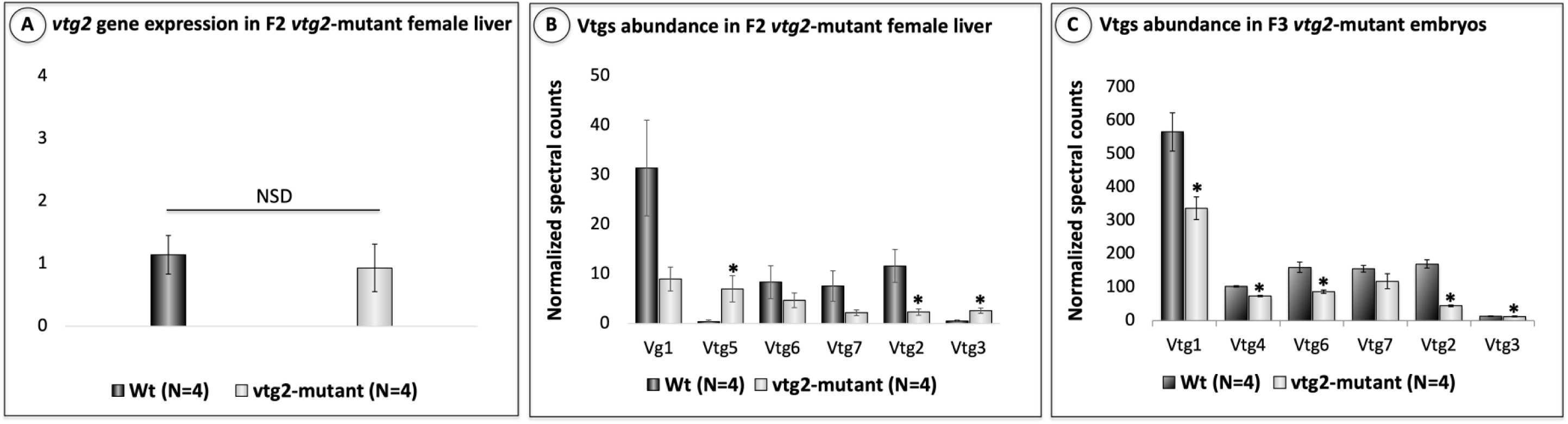
Relative quantification of *vtg2* gene expression in F2 *vtg2*-mutant zebrafish female liver and Vtg2 protein abundance in 1 hpf F3 *vtg2*-mutant embryos. A) Comparison of *vtg2* gene expression levels in F2 *vtg2*-mutant (Ht; *vtg2* -/+) and wild type (Wt) female liver in dark and light gray vertical bars, respectively. Vertical brackets indicate SEM. SYBR Green qPCR-2^-ΔΔCT^ mean relative quantification of gene expression normalized to the geometric mean expression of zebrafish elongation factor 1a (*eif1a*) and ribosomal protein L13a (*rpl13a*) was employed. Data were statistically analyzed using a Kruskal Wallis nonparametric test *p* < 0.05. **B)** Relative quantification of multiple vitellogenins by LC-MS/MS in F2 *vtg2*-mutant female (Ht; *vtg2* -/+) liver, and **C)** Relative quantification of multiple vitellogenins by LC-MS/MS in 1 hpf F3 *vtg2*-mutant embryos. Comparisons of mean normalized spectral counts for Vtg protein levels in Wt versus *vtg2*-mutants indicated by dark and light gray vertical bars, respectively. Vertical brackets indicate SEM. Statistically significant differences between group means detected by an independent samples Kruskal Wallis non-parametric test (p < 0.05) are indicated by asterisks above bars.

There were no significant differences between F2 *vtg2*-mutant (Ht; *vtg2* -/+) and Wt females in number of eggs per spawn (**Fig 5**). However, the fertilization rate, hatching rate and overall survival of F3 *vtg2*-mutant offspring were significantly different than seen in Wt fish (p < 0.05)(**Fig 5**). The fertilization rate of eggs from F2 *vtg2*-mutant females (36.95 ± 7.0 %) was substantially lower than those from Wt eggs (81.6 ± 5.22 %). About 22 % of the observed batches has a fertilization rate of absolute zero. About 34 % of the observed batches had <50 % while the remaining 45 % had ≥ 50 % fertilization rate. Hatching rate was significantly higher in F3 *vtg2*-mutant embryos in comparison to Wt embryos at 72 hpf (*vtg2*-mutant; 62.23 ± 5.6, Wt; 24.8 ± 8.54) and 96 hpf (*vtg2*-mutant; 97.19 ± 1.22, Wt; 70.6 ± 3.86, p < 0.05) (**Fig 5**). Notwithstanding, embryo and larval survival rates of F3 *vtg2*-mutant offspring were significantly less than of Wt offspring (p < 0.05), sharply decreasing to 29.3 ± 4.4 % at 24 hpf, to 18.2 ± 3.2 % on day 20 of the experimental period (**Fig 6**). About 40 % of the observed *vtg2*- mutant batches which mostly have demonstrated the yolk leaking disorder from the vitellin membrane had less than 10 % survival at 24 hpf, while 68 % of those has < 50 % survival ratio. Separate panels in **Fig 7** illustrate morphological disorders observed during development of F3 *vtg2*-mutant offspring in comparison to those from Wt females. Examples to yolk leakage due to vitellin membrane disintegrity and abnormal cell division are given in panels A and B. Phenotypic disorders mainly consist of pericardial and yolk sac/abdominal edema accompanied by spinal lordosis evidenced as curved or bent back deformities in *vtg2*-mutant embryo are shown in panels C and D.

**Fig 5.**
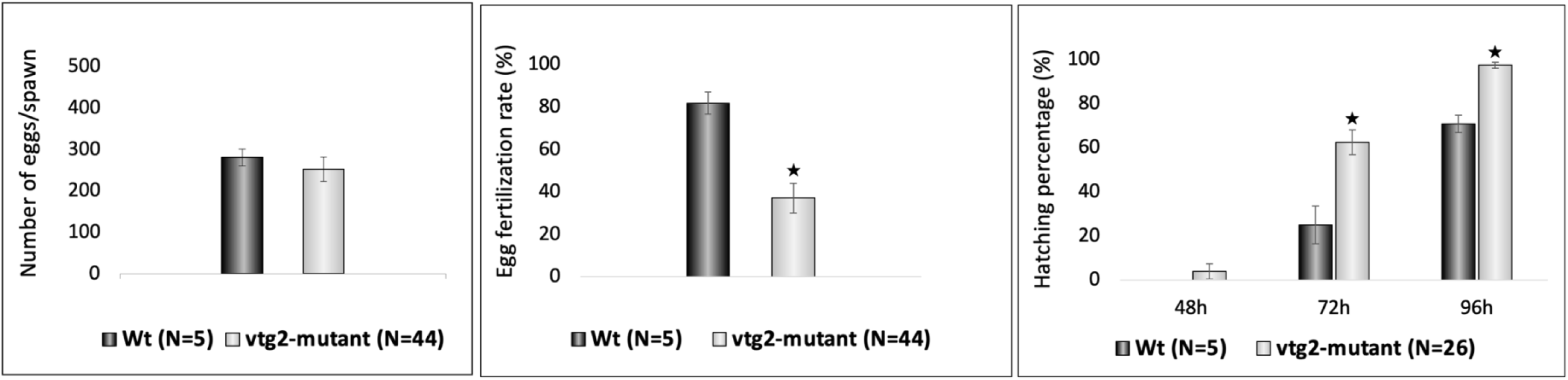
Phenotypic measurements of F2 *vtg2*-mutant females and their F3 progeny. Vertical bar graphs indicate mean values (± SEM) for measurements of **A)** number of eggs per spawn, **B)** egg fertilization rate and, **C)** embryo hatching percentages at 48, 72 and 96 hpf. Labels below the x-axes indicate the groups that were compared with the number of tested batches indicated in parentheses. Black stars, in all graphs, indicate mean values that are significantly different from corresponding Wt mean values based upon results of an independent samples t-test (p < 0.05).

**Fig 6.**
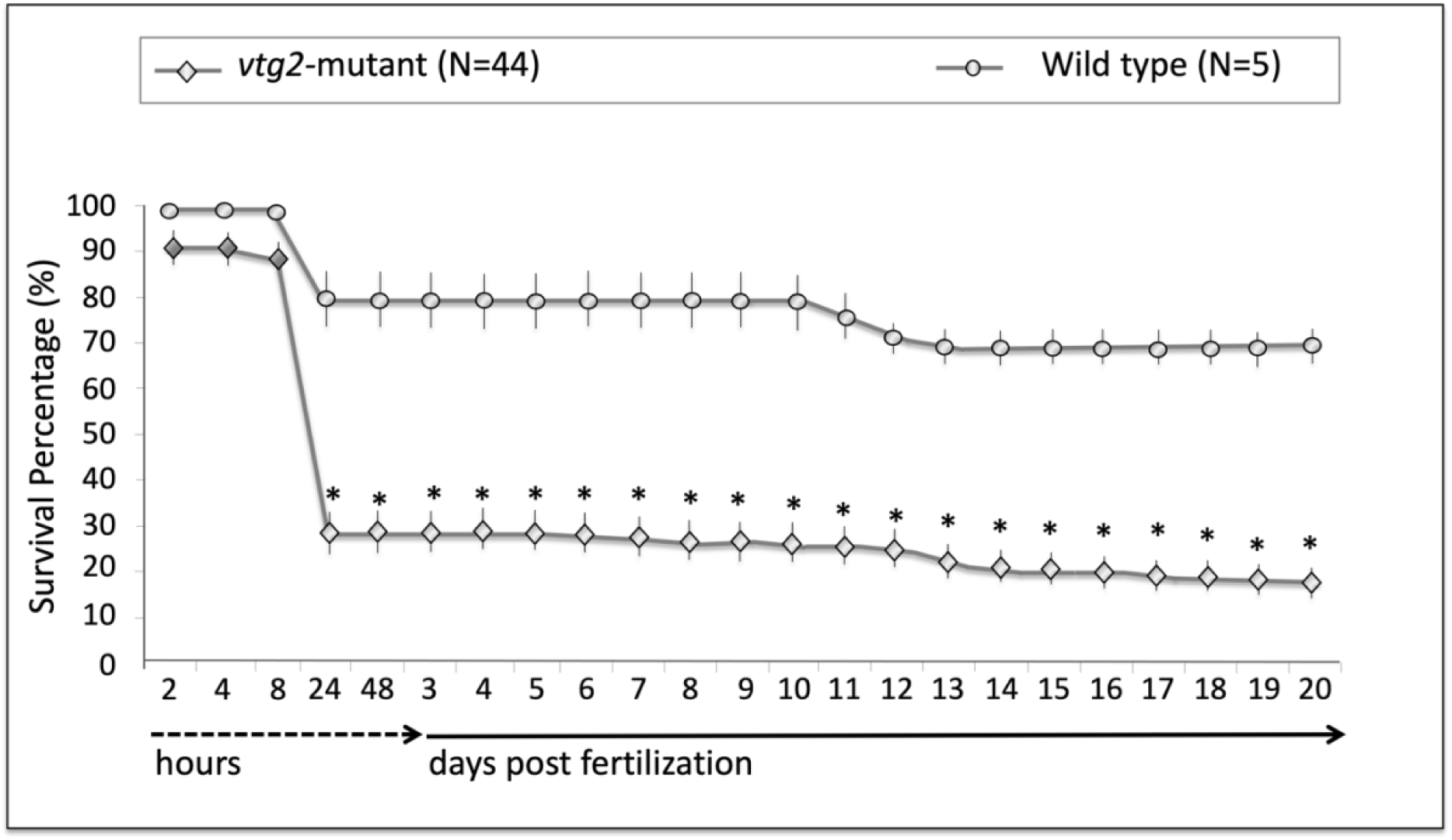
Comparisons of survival percentages for F3 *vtg2*-mutants versus wild type zebrafish offspring. Line plots represent mean survival percentages and numbers on the x-axis accompanied by dashed- and solid-lined arrows represent sampling times in hours or days post fertilization during the observation period. Mean survival percentages for *vtg2*-mutants and Wt embryos and larvae at each time point are indicated by diamonds and circles, respectively, and vertical lines indicate SEM. Asterisks indicate mean values that are significantly different from corresponding mean Wt values based upon results of an independent samples t-test (p < 0.05).

**Figure 7.**
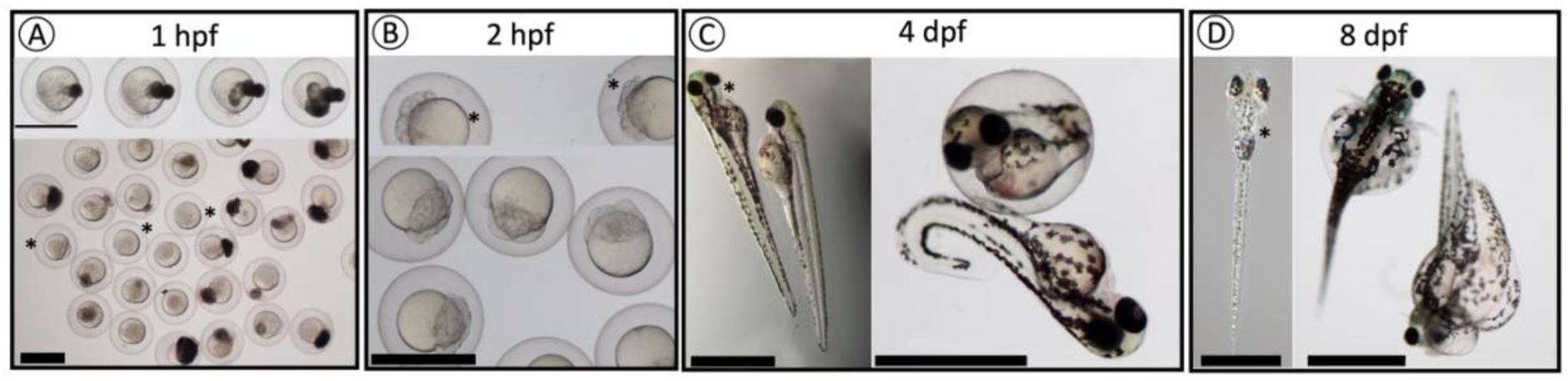
Observed phenotypes of F3 *vtg2*-mutant offspring compared to wild type offspring. **A)** *vtg2*-mutant embryos with leaking yolk at 1 hpf. **B)** *vtg2*-mutant embryos with abnormal cell division at 2 hpf. **C)** Wt versus *vtg2*-mutant embryos at 4 dpf. **D)** Wt versus *vtg2*-mutant embryos at 8 dpf. *Wt embryos are indicated by asterisks.

When extracts of newly fertilized F3 eggs from N = 4 F2 *vtg2*-mutant females and from N = 4 Wt females were compared in terms of their proteomic profiles via label-free LC-MS/MS, a total of N = 365 proteins was identified. Of these, N = 259 proteins were detected in at least four samples and showed a ≥ 1.0-fold difference in normalized spectral counts (N-SC) between groups (*vtg2*-mutant versus Wt), or were unique to a certain group, and, on this basis, were considered to be ‘differentially regulated’ (**S1 Table**). Accordingly, proteins which were detected with lower abundance in *vtg2*-mutant embryos in comparison to Wt embryos were further reported as “downregulated”, and those which were detected with higher abundance in *vtg2*-mutant embryos in comparison to Wt embryos were further reported as “upregulated” (**S1 Table**). The frequency distribution of differentially regulated proteins among 10 functional categories significantly differed (χ2, p < 0.05) between *vtg2*-mutant and Wt embryos (**Fig 8**). Frequencies of proteins related to cell cycle, division, growth and fate, protein degradation and synthesis inhibition, lectins and zona pellucida proteins were significantly higher, whereas frequencies of proteins related to protein synthesis, lipid metabolism and vitellogenins were significantly lower in *vtg2*-mutant embryos in comparison to Wt embryos. Individual proteins that were upregulated in *vtg2*-mutant embryos (N= 176, **Fig 8**) were mainly related to cell cycle, division, growth and fate (40 %) and lectins (23 %) with the remaining categorized proteins being related to energy metabolism (10 %), protein synthesis (6 %), protein degradation and synthesis inhibition (5%), zona pellucida proteins (5 %), redox/detox activities (3 %), immune functions (3 %), and Vtgs (2 %). Three percent of proteins which were upregulated in *vtg2*-mutant embryos were placed in the category of ‘others’. Individual proteins that were downregulated in *vtg2*-mutant embryos (N = 83, **Fig 8**) were mainly related to cell cycle, division, growth and fate (24 %), protein synthesis (22 %), energy metabolism (14 %), Vtgs (16%), and lectins (12 %), with the remaining categorized proteins being related to redox/detox activities (6 %), immune functions (4 %), and lipid metabolism (2 %).

**Fig 8.**
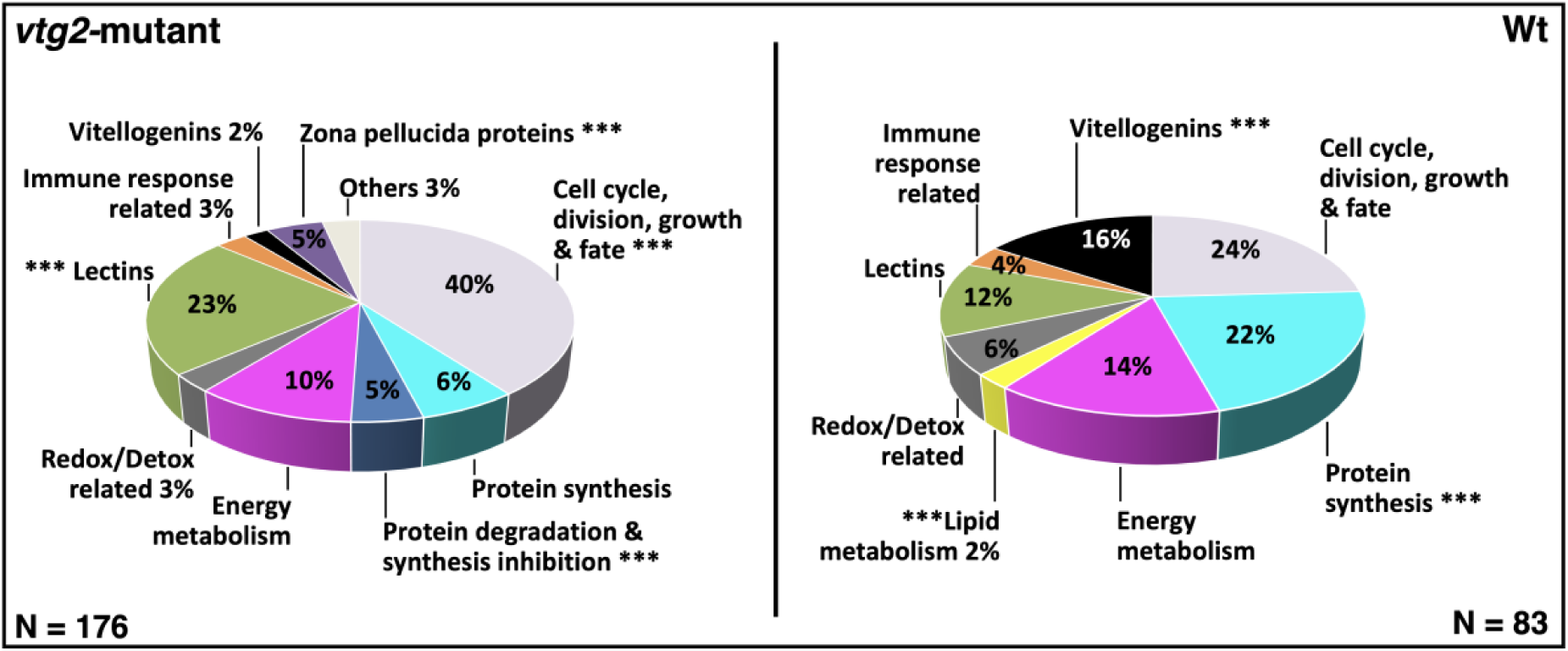
Percent distribution of DEPs. Distribution of differentially regulated proteins among functional categories. **Left Panel.** Proteins downregulated in *vtg2*-mutant offspring (upregulated in Wt, N = 83). **Right Panel.** Proteins upregulated in *vtg2*-mutant offspring (N = 176). Only proteins that were identified in > 4 biological samples and that exhibited a ≥ 1.0-fold difference in N-SC between groups (*vtg2*-mutant versus Wt), or proteins unique to a certain group, were included in this analysis. The overall distribution of differentially regulated proteins among the functional categories significantly differed between mutant and Wt eggs (χ2, p < 0.05). Asterisks indicate significant differences between different groups in the proportion of differentially regulated proteins within a functional category (χ2, p < 0.05). The corresponding Ensembl Protein IDs and associated gene, transcript and protein names, functional categories (shown above), regulation (upregulated or downregulated in mutant compared to Wt), and fold-difference in N-SC between *vtg2*-mutant and Wt eggs for proteins included in this analysis are given in **S1 Table**.

When the 259 differentially regulated proteins between *vtg2*-mutant and Wt embryos were submitted separately (*vtg2*-mutant downregulated (Wt); N=83, *vtg2*-mutant upregulated; N=176) to a functional protein association networks analysis using the Search Tool for the Retrieval of Interacting Genes/Proteins (STRING) and the zebrafish protein database, they resolved into networks with significantly and substantially greater numbers of known and predicted interactions between proteins than would be expected of the same size lists of proteins randomly chosen from the zebrafish database (**Fig 9**).

**Fig 9.**
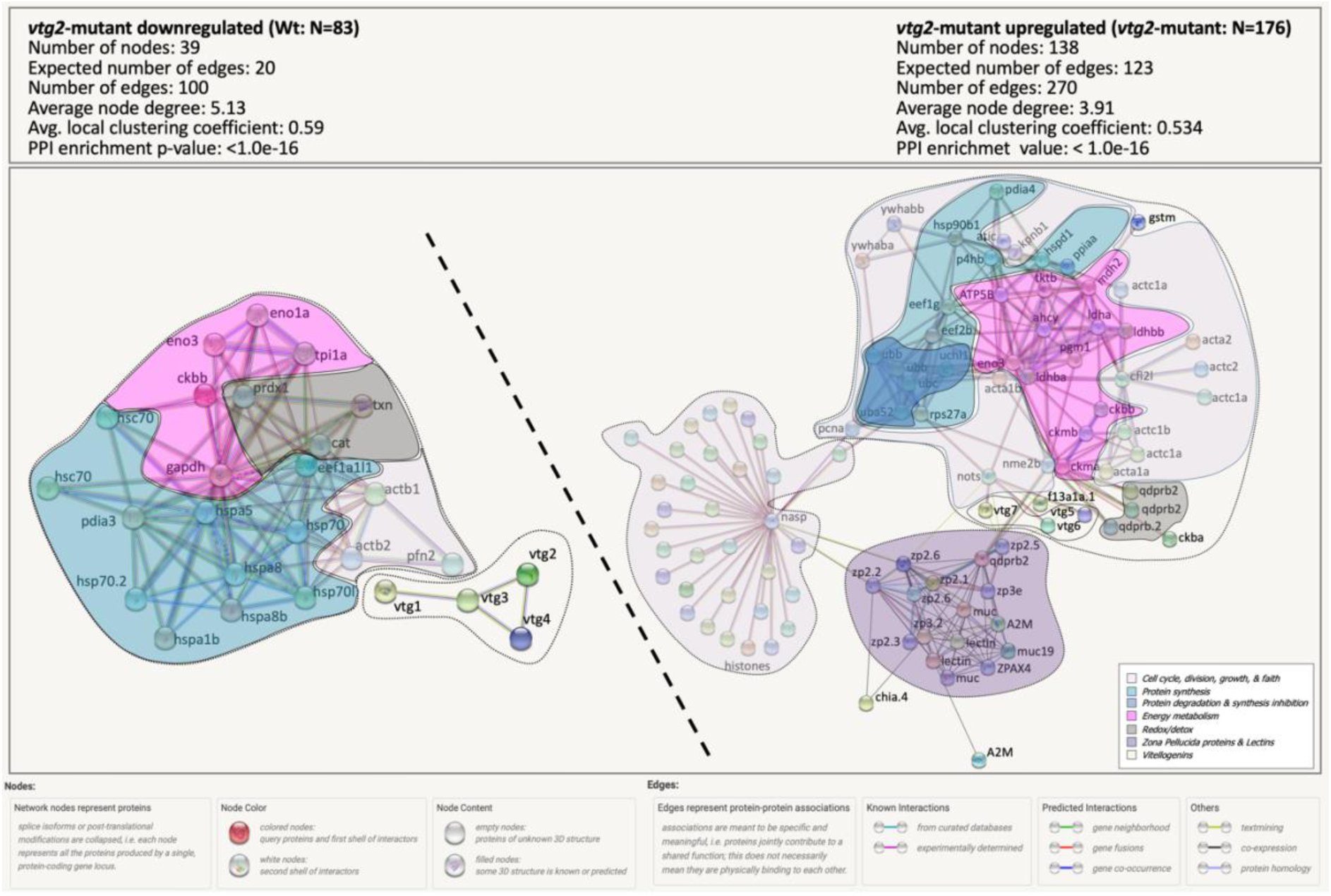
STRING Network Analysis of the differentially regulated proteins in *vtg2*-mutant eggs. A total of 83 proteins which were downregulated in *vtg2*-mutant eggs and 176 proteins which were upregulated in *vtg2*-mutant eggs were over-represented in specific biological pathways (**S2 Table**). Each network node (sphere) represents all proteins produced by a single, protein-coding gene locus (splice isoforms or post-translational modifications collapsed). Only nodes representing query proteins are shown. Nodes are named for the transcript(s) to which spectra were mapped; for full protein names, see **S1 Table**. Edges (colored lines) represent protein-protein associations meant to be specific and meaningful, *e.g.,* proteins jointly contribute to a shared function but do not necessarily physically interact. Protein clusters of similar functions are framed and highlighted with the same colors of functional categories in **Fig 8**. Color codes of these categories are given in the bottom left corner of the right panel. Model statistics are presented at the top left and at the top right of each panel for proteins down- and up-regulated in mutant eggs, respectively. Explanation of edge colors is given below panels. The subnetwork formed by proteins downregulated in mutant eggs is shown to the upper left above the diagonal dashed line, and the subnetwork formed by proteins upregulated in mutant eggs is shown to the lower right below the diagonal dashed line. Where possible, dashed lines encircle clusters of transcripts encoding interacting proteins involved in physiological processes distinct from other such clusters.

The network formed by proteins down-regulated in *vtg2*-mutant embryos (Wt; N= 83) is made up of a subnetwork of four Vtgs (Vtg1, Vtg2, Vtg3 and Vtg4) and another subnetwork made up proteins related to cellular homeostasis via cytoskeleton regulation, redox/detox activities, energy metabolism, and protein synthesis (protein folding and refolding activities). Cytoskeleton related proteins in this subnetwork are profilin (Pfn2), actin b1 (Actb1) and b2 (Actb2). Peroxiredoxin 1 (Prdx1), catalase (Cat) and thioredoxin (Txn) are proteins with Redox/detox functions within this subnetwork. Two forms of Enolases (Eno1a and Eno3) along with a creatinine kinase (Ckbb), a glyceraldehyde-3-phosphate dehydrogenase (Gapdh), and a triosephosphate isomerase 1a (Tpi1a) are proteins responsible of maintaining cellular energy homeostasis in this subnetwork. The eukaryotic translation elongation factor 1 alpha 1, like 1 (Eef1al1), along with two classes of heat shock proteins (Hspa, Hsp70), and a single protein disulfide-isomerase a3 (Pdia3) involved in protein folding and refolding of misfolded proteins were part of the protein synthesis cluster in this subnetwork (**Fig 9**. Left Panel, PPI network enrichment value p < 1.0e-16).

The network formed by proteins upregulated in *vtg2*-mutant embryos (*vtg2*-mutant; N = 176) is made up of three major interacting subnetworks. The first subnetwork is composed of proteins that are related to cellular homeostasis via cell cycle, division, growth and faith, protein synthesis, degradation and synthesis inhibition, energy hemostasis, and redox/detox activities. Among cell cycle, division, growth, and faith proteins are actins (Acta and Actc), cofilin 1 (Cfl1), karyopherin (importin) beta 1 (Kpnb1), 5-aminoimidazole-4-carboxamide ribonucleotide formyltransferase/IMP cyclohydrolase (Atic), two forms of tyrosine 3-monooxygenase/tryptophan 5-monooxygenase activation protein, beta polypeptide (Ywhaba and Ywhabb), proliferating cell nuclear antigen (Pcna), and NME/NM23 nucleoside diphosphate kinase 2b, tandem duplicate 2 (Nme2b). Protein synthesis cluster within this subnetwork includes translational proteins such as Eukaryotic translation elongation factors (Eef1g and Eef2b), Ribosomal protein S27a (Rps27) and folding/refolding related proteins such as Protein disulfide isomerase family A, member 4 (Pdia4), Heat shock protein 90, beta (grp94), member 1 (Hsp90b1), Heat shock 60 protein 1 (Hspd1), Peptidylprolyl isomerase Aa (cyclophilin A) (Ppiaa), and procollagen-proline, 2-oxoglutarate 4-dioxygenase (proline 4-hydroxylase), beta polypeptide (P4hb). Protein degradation and synthesis inhibition related proteins cluster includes ubiquitins which are mainly involved in response to unfolded/misfolded proteins (Uba52, Ubb, Ubc, and Uchl1), and an aspartic-type endopeptidase nothepsin (Nots). The central energy metabolism cluster of this subnetwork includes several creatine kinases (Ckba, Ckbb, Ckma, and Ckmb), lactate dehydrogenases (Ldha, Ldhab, and Ldhbb), phosphoglucomutase 1 (Pgm1), malate dehydrogenase 2 (Mdh2), enolase 3 (Eno3), transketolase b (Tktb), and ATP synthase F1 subunit beta (Atp5f1b). Positioned in the center of this cluster is an unrelated protein, adenosylhomocysteinase (Ahcy), an enzyme involved in breakdown of methionine to assist DNA methylation. A group of quinoid dihydropteridine reductase b2 (Qdprb2), proteins related to redox/detox activities, and a group of type I Vtgs (Vtg5, 6 and 7) are the remaining relatively smaller clusters spotted to be formed within this subnetwork. The second subnetwork formed by proteins upregulated in *vtg2*-mutant embryos exclusively includes histones and a single histone-binding protein, nuclear autoantigenic sperm protein (Nasp). And the third subnetwork includes zona pellucida proteins (ZPs), lectins and uncharacterized proteins recognized as mucins by the STRING algorithm (**Fig 9**. Right Panel, PPI network enrichment value p < 1.0e-16).

Functional enrichment analysis of proteins downregulated in *vtg2*-mutant embryos (**S2 Table**. Panel A) contributing to the identified protein-protein networks revealed the following Biological process GO-terms; Protein refolding, Chaperone cofactor-dependent protein refolding, Cellular response to unfolded protein, Protein folding, Glycolytic process, Generation of precursor metabolites and energy, Cellular response to stress, Nucleotide metabolic process, Regulation of vacuole fusion, non-autophagic, Carbohydrate metabolic process, Response to stress, Response to toxic substance. Molecular functions GO-terms for these proteins were; Protein folding chaperone, Misfolded protein binding, Nutrient reservoir activity, Unfolded protein binding, Heat shock protein binding, ATPase activity, Lipid transporter activity, Nucleoside-triphosphatase activity, 2,3-cyclic-nucleotide 3- phosphodiesterase activity, UDP-alpha-D-glucose:glucosyl-glycogenin alpha-D-glucosyltransferase activity, Glycogenin glucosyltransferase activity, Catalytic activity, Structural constituent of postsynaptic actin cytoskeleton, Phosphopyruvate hydratase activity, Low-density lipoprotein particle binding, Carbon-oxygen lyase activity, ATP binding, Nucleotide binding, Complement binding, Purine ribonucleoside triphosphate binding, Ubiquitin protein ligase binding, and Purine ribonucleotide binding. And KEGG pathways were Protein processing in endoplasmic reticulum, Spliceosome, Endocytosis, Carbon metabolism, MAPK signaling pathway, Glycolysis / Gluconeogenesis, Biosynthesis of amino acids, Salmonella infection, Starch and sucrose metabolism, Metabolic pathways.

Functional enrichment analysis of proteins upregulated in *vtg2*-mutant embryos (**S2 Table**. Panel B) contributing to the identified protein-protein networks revealed the following Biological process GO-terms; Nucleosome assembly, Chromosome organization, Protein-containing complex subunit organization, Mesenchyme migration, Cellular component assembly, Single fertilization, G protein-coupled receptor signaling pathway, Organelle organization, Phosphocreatine biosynthetic process, Tetrahydrobiopterin biosynthetic process, L-phenylalanine catabolic process, Cellular process, and Cell surface receptor signaling pathway. Molecular functions GO-terms for these proteins were Protein heterodimerization activity, Carbohydrate binding, Protein dimerization activity, Acrosin binding, Binding, G protein-coupled receptor activity, 6,7-dihydropteridine reductase activity, NADH binding, Protein tag, Creatine kinase activity, NADPH binding, Structural constituent of cytoskeleton, L-lactate dehydrogenase activity, Nutrient reservoir activity, DNA binding, Heterocyclic compound binding, and Organic cyclic compound binding. And KEGG pathways were Gap junction, Cysteine and methionine metabolism, Arginine and proline metabolism, Phagosome, Apoptosis, Tight junction, Glycolysis / Gluconeogenesis, Cardiac muscle contraction, and Pyruvate metabolism.

## 3. DISCUSSION

Our previous research demonstrated type I and type III Vtgs to have crucial functions during early embryo and larval development in zebrafish [9]. However, the role of type II Vtg remained to be revealed. The present research was undertaken to investigate the essentiality and functions of Vtg2 in zebrafish using complementary methodologies such as the CRISPR/Cas9 gene knock out approach, gene expression quantification using TaqMan qPCR and protein quantification using LC-MS/MS. Accordingly, an attempt to invalidate the *vtg2* gene has been made and subsequent effects on maternal reproductive physiology and offspring development and survival were assessed.

Efficacy of CRISPR/Cas9 in this study was ∼54 % and this was a better score in comparison to *vtg1*-KO experiment (20 %), where five genes were targeted concomitantly, but worse in comparison to the *vtg3*-KO experiment (80 %) where only a single gene was targeted [9]. The extremely low (0.25 %) mutation transmission was a serious limitation which complicated production of subsequent generations and pure mutation line in this study. However, the presence of heterozygous males at F0 and F1 generations allowed us to continue this study with single allele contribution of F2 generation females (Ht; *vtg2* -/+), which indeed enhanced the significance of our findings. Ultimately, F3 offspring batches that were subjected to proteomic and phenotyping analyses contained a mix of homozygous (Hm; *vtg2* -/-), heterozygous (Ht; *vtg2* -/+) and wild type (Wt; *vtg2* +/+) individuals with half maternal contribution because their female parent is a heterozygous with a single *vtg2* allele (Ht; *vtg2* -/+) (**Fig 3**). Significantly lower protein abundances of Vtg2 in F2 *vtg2*-mutant female liver and F3 generation offspring were indeed indicators of reduced functionality of *vtg2* in these heterozygous F2 females and their half contribution providing reduced *vtg2* supplies to the F3 generation offspring. And that in fact, the phenotypic and proteomic observations made from F3 generation offspring arise from reduced maternal Vtg2 contribution of Ht (*vtg2* -/+) F2 females to early development resources.

Even though the most part of it, including the receptor binding site, remained intact, by disturbing the structure of the LvH chain in *vtg2*-mutant genome via the creation of a large deletion (2811 bp, 564 aa) covering the C-terminal of the LvH polypeptide and the subsequent yolk protein domains (Pv, LvL, Bc, Ct), it is uncertain whether the mutant gene would be expressed and the protein would be synthetized, fold and transferred to the ovaries properly or not. However, qPCR and LC-MS/MS revealed limited function of *vtg2*/Vtg2 in *vtg2*-mutant individuals which is most probably resulting from the loss of a single allele in Ht; *vtg2* -/+ females. The non-significant lower gene expression of *vtg2* and significantly lower protein abundance in the liver of mutant F2 females substantiate the fact that these females are Ht; *vtg2* -/+ carrying a single *vtg2* allele. Transcripts of other type of *vtg* genes in mutant female liver were not quantified in this study. However, the Vtg2 protein levels were still detectable but significantly lower in both F2 *vtg2*-mutant female liver and in F3 *vtg2*- mutant offspring in comparison to Wt offspring. These results corroborate the reduced contribution of the Ht; *vtg2* -/+ females to Vtg2 stockpile in F3 *vtg2*-mutant offspring.

The ‘genetic compensation’ phenomenon which has been observed in *vtg1*- and *vtg3*-KO experiments has also been encountered in the form of significant increase in Vtg5 and Vtg3 abundance in *vtg2*-mutant female liver in this study. However, this increase did not reflect in the Vtgs profiles of the F3 *vtg2*-mutant offspring. Protein abundance of Vtg1, 4, 6 and 3 were found to be significantly lower in F3 *vtg2*-mutant embryos. The reason to this decrease is unknown and worth further investigation. *vtg2*-mutation did not influence the number of eggs per clutch that F2 *vtg2*-mutant females spawned, however, fertilization rate of *vtg2*-mutant eggs was only ∼37 %, less than half of that was observed in Wt eggs. Moreover, 20 % of the observed F3 vtg2-mutant batches had absolute zero fertilization. Similar results were found in eggs from Hm *vtg3*-KO females (35.5 ± 7.7 %) in our previous study [9]. In both studies, fertilization rate was estimated conservatively, based on numbers of viable embryos showing normal cell division and subsequently developing to ∼24 hpf. It is uncertain whether the high mortality of *vtg2*-mutant eggs at 24 hpf resulted from a failure to be fertilized or from defects in early development involving zygotes that fail to initiate cell division or that briefly undergo abnormal cell divisions and then die. The mechanism(s) whereby Vtg2 deficiency impairs fertility of zebrafish are unknown, however, Vtg2 is clearly a major contributor to fertility and/or early development in zebrafish. Interestingly, eggs from F2 *vtg2*-mutant females exhibited significantly higher embryo hatching rates at 72 and 96 hpf (*p* < 0.05) in comparison to eggs from wild type females. This unique phenomenon might well be related to over- representation/expression of ZP proteins and vitellin membrane deformities (*see below*). The continuous mortality of F3 *vtg2*-mutant embryos after 24 hpf leading to only ∼18 % survival at 20 dpf, suggests that Vtg2 also contributes to late embryonic and larval development, as suggested in several prior studies. Embryo and larval survival rates were severely diminished by *vtg*2 disturbance. The substantial losses of *vtg2*-mutant eggs began early and less than 30% survived to 24 hps. Survival of embryos emanating from *vtg2*-mutant females decreased drastically to less than 30 % at 24 hpf and further to ∼16 % thereafter, becoming significantly less than survival of Wt embryos during the experimental period. It is also important to note that F3 *vtg2*-mutant batches theoretically contained 50 % *vtg2* -/+, 25 % *vtg2* -/-, and 25 % *vtg2* +/+, but all received 50 % reduced Vtg2 supplies from their Ht; *vtg2* -/+ female parent.

Like the *vtg1- and vtg3*-KO zebrafish larvae, *vtg2*-mutant survivor offspring exhibited major phenotypic disorders, mainly pericardial and yolk sac/abdominal edemas and spinal lordosis associated with curved or arched back deformities. However, those which were lost at earlier stages exhibited clear defects evidenced by leakage of the yolk throughout the vitellin membrane. *vtg2*-mutant embryos unconditionally exhibited serious and lethal developmental abnormalities. Similarities in observed phenotypes following the loss of type I, II and III Vtgs in zebrafish might imply some commonalities in functions of these three different types of Vtgs, however, no certain evidence to that was possible to identify through our studies.

Proteomic profiles of *vtg2*-mutant embryos were significantly different from those of Wt embryos. The results of frequency distribution analysis were in agreement with those of STRING network enrichment analyses, attributing a far greater proportion of proteins to GO terms related to cell cycle, division, growth and fate, ZP proteins and Lectins. Most of the cell cycle, division, growth and fate related proteins were H4 histones and proteins with H4 histone domains. Histones are core proteins of the chromatin, a fundamental unit of nucleosome, composed of DNA wrapped around these core proteins. The maturing Xenopus oocyte and laid egg have been reported to contain an accumulated store of nonchromatinized, predeposition histones complexed with chaperones [10]. Studies from Bao et al., [11] identify the NASP, which is in the center of the Histone H4 cluster in our STRING protein-protein network formed by proteins upregulated in *vtg2*-mutant embryos, as a major H3–H4 chaperone in histone homeostasis maintenance. The store of histones in the egg has been hypothesized to be required for the rapid cell cycles in the developing frog prior to the midblastula transition [10]. Massive dead tolls at first cell division cycles during early development in *vtg2*-mutant embryos, and concurrent overrepresentation and upregulation of histone 4 proteins in our study suggest a potential adaptive response to cope with developmental deficiencies caused by the genetic disturbance of *vtg2* in the female parent causing reduced Vtg supplies in the F3 offspring.

In addition, histones are subject to a wide range of post-translational modifications, such as acetylation, phosphorylation, and ubiquitination, that play critical roles during early embryo development [10]. Histone acetylation is a regulatory modification that occurs predominantly at the level of lysines in the aminoterminal tail of histones 3 and 4. It plays an important role in gene expression by altering the accessibility of DNA to proteins, such as transcription factors. In the majority of cases, genes that are undergoing transcription are hyperacetylated, and those that are repressed are deacetylated [12, 13, 14]. Earlier studies explored changes in histone modifications and DNA methylations in relation to zona pellucida removal in mice, but only DNA methylations were reported to be declined significantly at 2-4 cell stages of early embryonic development [12]. Despite the lack of evidence to a link between the presence of zona pellucida and histone modifications, the high abundance and distribution of H4 histones and the parallel upregulation of zona pellucida proteins in 1 hpf *vtg2*- mutant embryos is curious and begs further investigation.

Zona radiata or zona pellucida, vitellin envelope, perivitellin envelope, is a relatively thick extracellular matrix or coat, which surrounds both invertebrate and vertebrate eggs [15]. In fish, the egg envelope is different in structure and has disparate function in comparison to mammals. The eggshell in teleost eggs consists of an extremely thin, high-density outer layer and a thick, low-density inner layer, which makes up most of the structure [16]. Studies on localization of sea bream ZP proteins in the ZP matrix indicated the ZPX to be exclusively situated in the innermost layer, the ZPB and ZPA (ZP1 and 2) to contribute to the central portion, while the ZPC (ZP3) to be the solely constituent of the outer layer of the vitellin membrane [17]. According to this, upregulation of ZP1, 2, 3 in our study is somehow related to premature disintegration of the central and outmost layers of the vitellin membrane in *vtg2*-mutant embryos. Upregulation of ZP 2 and 3 proteins was also reported in poor quality zebrafish eggs [18], however, the yolk leakage phenomenon has been for the first time observed in *vtg2*-mutant embryos and can be considered as evidence to these malformations.

According to Hara et al. [16], the timing of eggshell formation differs by layer, and in general, the outer layer formation begins by the perinucleolar oocyte stage while the formation of the inner layer coincides with previtellogenic oocyte development. Thus, by the time of inner layer formation, the oocyte is being filled with yolk proteins derived from vitellogenins and very-low density lipoproteins carried to the oocyte by vitellogenins. Depressing of the *vtg2* function in our study might somehow have disturbed this natural sequence of events that led to disintegration of the vitellin membrane in *vtg2*- mutant embryos under insufficient Vtg provisions.

Similar to the protein profiles obtained from poor quality zebrafish eggs (1 hpf), sea urchin egg lectin (SUEL)-type and Ca2^+^-dependent (C-type) lectins were significantly elevated in number and abundance in *vtg2*-mutant embryos. Moreover, like good quality zebrafish eggs (1 hpf), fish egg lectin (FEL)-like proteins were higher in numbers and abundance in wild type embryos when compared to *vtg2*-mutant embryos in our study. C-type lectins are known to be localized at the cortical granules of fish eggs. From here they are discharged at fertilization into the perivitelline space, where they function in chorion hardening of the egg and establishing the block to polyspermy [19]. Overrepresentation and expression of these lectins in *vtg2*-mutant embryos indicate contribution of the lectins into the adaptive response to maintain the integrity of the vitellin envelop.

In contrast to proteomic profiles of *vtg1*- and *vtg3*-KO embryos multifunctional endoplasmic reticulum resident redox chaperones, [20] such as heat shock proteins (*e.g.,* Hsp70 kDa) and protein disulfide-isomerase (Pdia3) were downregulated in *vtg2*-mutant embryos. These proteins are responsible for the correct folding and maintenance of the native protein conformation, and therefore, serve as a staminal cellular defense against general protein misfolding [21]. Downregulation of these proteins in *vtg2*-mutant embryos might indicate a failure of the cell to cope with ER stress conditions, leading to erroneous protein conformations and vitellin membrane degradation. Higher representation of proteins related to protein degradation and synthesis inhibition mechanisms in *vtg2*-mutant embryos in addition to biological processes and molecular functions revealed by functional enrichment and KEGG pathway analyses of proteins downregulated in *vtg2*-mutant embryos provide additional evidence to this conjecture.

In conclusion, this study provides a unique overview on essentiality *vtg2* in early embryonic development and procure empirical evidence to signatures of molecular changes and subsequent cellular dysfunctions caused by its partial genetic disturbance in zebrafish. These signatures of impairments in the cellular functions were possible to observe as early as at 1 hpf and were in accordance with phenotypical observations. Further research is needed if the complete disturbance effects in homozygous fish to be pursued, however, findings of this study state unequivocally that the Vtg2 has critical requisite functions for oogenesis and embryonic and larval development in zebrafish. Findings of this study also allowed the discovery of common molecular signatures between *vtg2*-mutant, vtg*1*- and *vtg3*-KO embryos and poor quality zebrafish eggs and the establishment of the overall portrait on disparate functions of multiple Vtgs in zebrafish, providing an excellent guide to study involved mechanisms in other oviparous vertebrates.

## 4. MATERIAL AND METHODS

### 4.1. Animal care

Zebrafish of the Tübingen strain originally emanating from the Nüsslein-Volhard Laboratory (Germany) were obtained from our zebrafish facility (INRA UR1037 LPGP, Rennes, France). The fish were ∼15 months of age and of average length ∼5.0 cm and average weight ∼1.4 g. The zebrafish were housed under standard conditions of photoperiod (14 hours light and 10 hours dark) and temperature (28 °C) in 10 L aquaria, and were fed three times a day *ad libidum* with a commercial diet (GEMMA, Skretting, Wincham, Northwich, UK). Females were bred at weekly intervals to obtain egg batches for CRISPR sgRNA microinjection (MI). The night before spawning, paired males and females bred from different parents were separated by an opaque divider in individual aquaria equipped with marbles at the bottom as the spawning substrate. The divider was removed in the morning, with the fish left undisturbed to spawn. Egg batches in majority containing intact, clean looking, well defined, activated eggs at the 1-cell stage were immediately transferred to microinjection facilities.

### 4.2. Single guide RNA (sgRNA) design, synthesis and microinjection

The *vtg2* genomic sequence was submitted to online available target designer tool at http://zifit.partners.org/ZiFiT/ChoiceMenu.aspx [22, 23] and of the proposed candidates, three regions located on exons 4, 20 (corresponding to the LvH yolk protein domain) and 32 (corresponding to the βc yolk protein domain) were chosen as target sites for this experiment. A schematic representation of the general strategy followed for CRISPR target design is presented in **Fig 1**. Forward and reverse oligonucleotides matching the chosen target sequences (given in **S3 Table)** were annealed and ligated to the pDR274 expression vector (Addgene). The vector was subsequently linearized by the DraI restriction digestion enzyme (Promega) and *in vitro* transcribed using mMessage mMachine T7 Transcription Kit (Ambion) according instructions from the manufacturer. The pCS2-nCas9n plasmid (Addgene Plasmid 47929) was digested with Not I restriction digestion enzyme (Promega) and transcribed using mMessage mMachine SP6 Transcription Kit (Ambion) according instructions from the manufacturer. The sgRNA concentration was measured on a Nanodrop 1000 Spectrophotometer (Thermo Scientific, USA) and integrity was tested before use using an Agilent RNA 6000 Nano Kit (Agilent) on an Agilent 2100 Bioanalyzer.

Approximately 100 eggs per batch were injected with sgRNA mix containing sgRNAs for three target sites at ∼30 ng/ul each and nCas9n RNA at ∼200 ng/ul at the one-cell stage. A total of 120 pg sgRNA mix and ∼800 pg Cas9 RNA was injected per embryo. Injected embryos were kept in 100 mm Petri dishes filled with embryo medium (17.1 mM NaCl, 0.4 mM KCl, 0.65 mM MgSO_4_, 0.27 mM CaCl_2_, 0.01 mg/L methylene blue) to assess microinjection efficiency, embryo survival and development post injection.

### 4.3. Genotyping by conventional PCR

As representatives of their generation, ten embryos were sampled randomly and gDNA was extracted individually and used as a template in targeted conventional PCR reactions to screen for introduced mutations in the targeted *vtg* genes. For this purpose, embryos surviving for 24 h post-injection were incubated in 100 µl of 5 % Chelex® 100 Molecular Biology Grade Resin (BioRad) and 50 µl of Proteinase K Solution (20 mg/ml, Ambion) initially for 2h at 55 °C and subsequently for 10 min 99 °C with constant agitation at 12000 rpm. Extracts were then centrifuged at 5000 xg for 10 minutes and supernatant containing gDNA was transferred into new tubes and stored at -20 °C until use.

To evaluate generational transfer of introduced mutations, genotyping of ∼2 month-old descendants was conducted after extraction of gDNA from fin-clips. For this purpose, fish were anaesthetized in 2-phenoxyethanol (0.5 ml/L) and part of their caudal fin was excised with a sterile scalpel. Genomic DNA from fin tissues were then extracted using Chelex 5 % as described above.

One µl (∼ 100 ng) of extracted gDNA was used in 20 µl PCR reactions using AccuPrime^TM^ Taq DNA Polymerase, High Fidelity (Invitrogen) and 10x AccuPrime^TM^ PCR Buffer II in combination with gene specific primers (at 10 µM each) anchoring target sites on the genomic sequence of targeted genes (**Fig 1**, **S3 Table**). PCR cycling conditions were as follows; 1 cycle of initial denaturation at 94 °C for 2 min, 35 cycles of denaturation at 94 °C for 15–30 sec, annealing at 52–64 °C for 15–30 sec and extension at 68 °C for 1 min per kb plus 1 cycle of final extension at 68 °C for 5 min. Non-purified PCR products or gel purified DNA were sequenced using gene specific primers indicated in **S3 Table** by the Eurofins Genomics sequencing service (https://www.eurofinsgenomics.eu/). Obtained sequences were aligned to corresponding zebrafish genomic sequence using Clustal Omega [24] for characterization and localization of introduced mutations, and then were blasted against all sequences available online using NCBI nucleotide Blast (Blastn) [25] for confirmation of the consistency, accuracy and type of the mutations created at the target sites.

### 4.4. Generation of zebrafish mutant lines

The initial strategy to produce the pure mutant line involved crossing of adults carrying the introduced mutation from each generation to produce the next generation until the pure line is obtained. In the case of our study, due to limited availability of survivor mutants, slight changes to this strategy have been applied. A single heterozygous (Ht; *vtg2*+/-) male with the mutation on a single allele was outcrossed with wild type (Wt; *vtg2*+/+) female with no genomic disturbance to produce the F1 generation. Embryos from F1 generation were genotyped to confirm mutation transfer and remaining embryos were raised to adulthood. F1 descendants were screened again at ∼2 months of age and, since mutation transmission was possible to detect in two males only, these Ht males were crossed with Wt females to produce the F2 generation. Following the same genotyping strategy, F2 Ht males were crossed with Ht females to produce the F3 generation. Low number of fish with mutations at each generation proved our attempt to produce the pure *vtg2*-KO lines unfeasible. Females from F2 generation (N=4) and from non-related wild type females (N=4) were spawned and then euthanized with a lethal dose of 2-phenoxyethanol (0.5 ml/L) to collect liver tissue for *vtg2* gene expression and protein quantifications. F3 *vtg2*-mutant and wild type embryo samples (N=4 each), from the same females, were collected at 1 hpf and snap frozen to assess label free protein profiling and quantifications via LC-MS/MS.

### 4.5. Phenotypic observations

For phenotypic observations Wt and F2 *vtg2*-mutant couples (N=5 each) were spawned from 1 to 8 times, respectively, and embryonic development, survival rate, hatching rate, and larval development were subsequently observed until 20 dps. Fecundity and fertilization data was collected from a total of 5 and 44 batches from Wt and F2 *vtg2*-mutant couples, respectively. Survival and hatching ratio data was collected from offspring (F3) of the same couples. Fecundity (number of eggs per spawn) was recorded immediately after spawning and collected eggs were incubated in 100 mm Petri dishes filled with embryo medium (17.1 mM NaCl, 0.4 mM KCl, 0.65 mM MgSO_4_, 0.27 mM CaCl_2_, 0.01 mg/L methylene blue) to assess embryonic development and phenotyping parameters. Hatching ratio was calculated based on the number of surviving embryos at 24 h and only spawns with > 5 % survival rates were considered, therefore, this data was collected from 5 of Wt and 26 of *vtg2*-mutant batches, respectively. Incubated eggs/embryos were periodically observed at the early blastula (∼256 cell) stage (∼2-3 h post spawning hps), at mid-blastula transition stage (∼4 hps), at the shield to 75% epiboly stages (∼8 hps), at the early pharyngula stage (∼24 hps), and during the hatching period at 48, 72 hps and 96 hps (long-pec to early larvae stages) following standard developmental staging [26]. Fertilization rate was calculated based on viable embryos showing normal cell division and subsequent development to ∼24 hpf since zygotes failing to initiate cell division, and embryos showing asymmetrical cell cleavage or early developmental arrest were dead by then. The number of surviving eggs/embryos was recorded, those not surviving were removed and the number of abnormal embryos was recorded at each observation point. Hatched embryos were transferred into larger volume containers (1 L) filled with standard 28°C culture water and were fed *ad libitum* with artemia and GEMMA weaning diet mix after yolk sac absorption (at around 10 dpf). At the time of feeding, larvae were also observed for motor and feeding activities. Observations were made daily up to 20 dpf.

### 4.6. Quantitative real time PCR

Total RNA was extracted from frozen liver using TriReagent (SIGMA) and cDNA was synthesized using SuperScript III reverse transcriptase (Invitrogen, USA) from 1 µg of total RNA according to the manufacturer’s instructions. Relative expression levels for zebrafish *vtg2* in F2 *vtg2*- mutant and wild type female liver were measured using SYBR GREEN qPCR Master Mix (SYBR Green Master Mix kit; Applied Biosystems) as indicated by the manufacturer in a total volume of 10 µl, containing RT products diluted at 1:100 and 400 nM of each primer in order to obtain PCR efficiency between 95 and 100 %. Sequences of primers used in this experiment are given in S1 Table. The RT-qPCR cycling protocol included 3 min initial denaturation at 95 °C followed by 40 cycles of 95 °C for 3 sec and 60 °C for 30 sec on a StepOnePlus thermocycler (Applied Biosystem). The relative abundance of target cDNA within a sample set was calculated from a serially diluted cDNA pool (standard curve) using Applied Biosystem StepOne V.2.0 software. Similarly, the 2^-ΔΔCT^ mean relative quantification of gene expression method with the mean expression value of zebrafish elongation factor 1a (*eif1a*) and ribosomal protein 13a (*rpl13a*) were employed as housekeeping genes in this study. Primer sequences and properties for these genes are also given in **S3 Table**. Obtained data was subjected to independent samples Kruskal-Wallis nonparametric test (p < 0.05) (IBM SPSS Statistics Version 19.0.0, Armonk, NY).

### 4.7. Liquid Chromatography Tandem Mass Spectrometry

Liver tissues from F2 *vtg2*-mutant and Wt females, and 1 hpf F3 embryos (N=5 batches each) from the same females were subject to protein extraction and sample processing for LC-MS/MS as described by [18] with slight changes. Concentration of extracted proteins was determined by Bradford Assay [27] (Bio-Rad, Marnes-la-Coquette, France). Samples of extracts were mixed with sample buffer and DTT and denatured at 70 °C for 10 min before being subjected to SDS-PAGE (60 µg protein/sample lane on NuPAGE 12% Bis-Tris Gel) for ∼4 minutes at 200V 218 mA (∼44 W) in NuPAGE™ MES SDS Running Buffer (Thermo Fisher Scientific, Illkirch-Graffenstaden, France). When protein samples had completely penetrated the stacking gel electropohoresis was stopped and gels were briefly rinsed in MilliQ ultrapure water (Millipore S.A.S., Alsace, France) and then incubated in fixation solution containing 30 % EtOH / 10 % acetic acid / 60 % MilliQ water for 15 min in order to fix proteins on the gel. Gels were then washed in MilliQ water and stained in EZBlue™ Gel Staining Reagent (Sigma-Aldrich, Saint-Quentin Fallavier, France) for 2 h, and de-stained in MilliQ overnight. Subsequently, protein bands were excised from the gel and processed for in-gel tryptic digestion and peptide extraction as indicated by [18]. Peptide extracts were resolubilized in 30 μl of 95 % H_2_O : 5 % formic acid and diluted 10 times before being injected to the LC-MS/MS system.

The LC-MS/MS system consisted of a nanoflow high-performance liquid chromatography (HPLC) (LC Packings Ultimate 3000, Thermo Fisher Scientific, Courtaboeuf, France) connected to a hybrid LTQ-OrbiTrap XL spectrometer (Thermo Fisher Scientific) equipped with a nanoelectrospray ion source (New Objective), as previously described [28, 18]. Obtained MS/MS spectra were searched against a target-decoy concatenated database created from the zebrafish Ensembl proteome database (Danio rerio_Zv9, March 2015) using Mascot (Matrix Science). Protein identification, quantification by spectral counts, and spectral count normalization were conducted as described by Yilmaz et al. [18]. The mass spectrometry proteomics data have been deposited to the ProteomeXchange Consortium [29] via the PRIDE [30] partner repository with the dataset identifier PXD042791. To detect significant differences in protein abundances based on group mean N-SC values (Wt vs *vtg2*-mutant) an independent samples Kruskal-Wallis nonparametric test (p < 0.05) followed by Benjamini-Hochberg correction for multiple tests (p < 0.25) was used (IBM SPSS Statistics Version 19.0.0, Armonk, NY).

### 4.8. Protein identification, quantification, annotation and statistics

A list with a total of 365 proteins identified in Wt and 1 hpf F3 *vtg2*-mutant embryos was filtered to exclude those detected in less than 4 biological samples and have less than 1.0 fold difference in abundance and the remaining proteins (N = 259) were identified using online available functional annotation tools such as Gene Ontology, Kyoto Encyclopedia of Genes and Genomes and Database for Annotation, Visualization and Integrated Discovery (DAVID) [31, 32, 33, 34]. These proteins were then classified into thirteen arbitrarily chosen functional categories that would account for > 90% of the proteins as originally suggested by Yilmaz et al. [18]. Chi square analysis with significance level of (p < 0.05) was used to detect differences between groups in the distribution of differentially regulated proteins among functional categories. Proteins which were differentially regulated in 1 hpf F3 *vtg2*- mutant embryos were subjected to the additional analysis of protein-protein interaction networks [35] and the functional enrichment within identified interaction networks separately using the STRING Network search tool available from the STRING Consortium online at http://string-db.org/, with the data settings Confidence: Medium (0.40), Max Number of Interactions to Show: None/query proteins only. For the Only statistically significant enrichment results from this analysis (p < 0.05) are reported.

## ACKNOWLEDGEMENTS

The authors thank Dr. Amaury Herpin for his advice on guide RNA selection, Dr. Rolf Brudvik Edvardsen on his advice on genotyping strategies, Amélie Patinote and the INRAE ISC-LPGP staff for fish breeding. Protim Core Facility was supported by structural grants from Infrastructures en Biologie Santé et Agronomie (IBiSA), Conseil Régional de Bretagne (through Biogenouest), European Regional Development Fund, Rennes Métropole and University of Rennes (through Biosit). This study was supported by FP7 People 2013-Marie-Curie Actions-Intra-European Fellowships (FP7-PEOPLE-2013-MC-IEF), Grant/Award Number: 626272; Region Bretagne in France-(SAD-2013), Grant/Award Number: 13009218; and Agence Nationale de Recherches, Grant/Award Number: ANR-13-BSV7- 0015.

## 5. SUPPORTING INFORMATION CAPTIONS

**S1 Fig. A)** Position of target sites and screening primers on *vtg2* gDNA. Introns are indicated in cyan, exons are indicated in plain text, screening primers are indicated in underlined bold text, target sites are indicated in magenta highlighted text along with corresponding labels to the right of the page, respectively. 21Fw, target1 forward primer; 21Rv, target1 reverse primer; 22Fw, target2 forward primer; 22Rv, target2 reverse primer; 23Fw, target3 forward primer; 23Rv, target3 reverse primer. sg21, sg22, and sg23: single guide RNAs (sgRNAs) for target sites 1, 2, and 3 for vtg2, respectively. Introduced mutation is indicated by strikethrough text. **B)** Position of target sites and screening primers on vtg2 cDNA. Sequence regions corresponding to each yolk protein domain are indicated with the following color highlighted text pointed out with brackets; LvH: plain text PV: orange, LvL: green, Bc: red, Ct: purple. Target sites are indicated in magenta highlighted text along with corresponding labels to the right of the page in front of brackets. sg21, sg22, and sg23: single guide RNAs (sgRNAs) for target sites 1, 2, and 3 for vtg2, respectively. Introduced mutation is indicated by strikethrough text. **C)** Predicted translational frames after deletion on *vtg2*. 5’3’Frame 3 is considered as the potential translation following the introduced mutation. **D)** Position of the introduced mutation on Vtg2 peptide sequence. Regions corresponding on each yolk protein domain are indicated in the following color highlighted text; LvH: plain text, Pv: orange, LvL: green, Bc: red, Ct: purple. Introduced mutation is indicated by strikethrough text.

**S1 Table.** Proteins differentially regulated in the *vtg2*-mutant eggs. List of the 259 proteins were considered to be differentially regulated between wild type (Wt) and vtg2-mutant eggs whose distribution among various functional categories is illustrated in **Fig 8**. These include proteins downregulated in vtg2-mutant eggs (Wt: N=83) and those upregulated in vtg2-mutant eggs (vtg2- mutant: N=176) with N-SC ≥1-fold relative to values for Wt eggs. Only proteins that were detected in at least 4 experimental individuals were considered in this study. For each protein, the Ensembl Protein ID and associated gene, transcript and protein name, functional category, relative abundance (down- or upregulated in vtg2-mutant), fold-difference in N-SC between groups (if available), and significance of differences (p < 0.05 and Benjamini-Hochberg corrected p < 0.25) is shown. Colour shading corresponds to that used to designate functional categories in **Fig 8**.

**S2 Table.** Functional enrichment analysis of differentially abundant proteins in STRING protein-protein interactions networks.

**S3 Table**. Targets and primers utilized in this study. Target oligo and screening primer names are given according to **Fig 1**. CRISPR recognition NGG motifs are highlighted by bold typeface on sequences. Position of primers and target sites on vitellogenin (Vtg) yolk protein (YP) domains are given on the far right columns.

